# Reproductive history and cognitive aging: Interactive effects of children and grandchildren in a life history framework

**DOI:** 10.64898/2026.05.26.727926

**Authors:** Oscar R. Sánchez, Ana María Salazar Montes, Olga Lucía Pedraza, Juan David Leongómez

## Abstract

Cognitive decline, particularly Alzheimer’s disease, represents a significant global health concern. While traditional risk factors are well documented, an evolutionary perspective grounded in life history theory offers critical insights into the intergenerational dynamics influencing cognitive aging. This study empirically analyzed the relationship between reproductive history and late-life cognitive performance in women, as measured by the Montreal Cognitive Assessment Colombian version (MOCA-Col). Specifically, we examined the moderating role of the number of grandchildren on the association between parity and cognitive performance. Using an observational, cross-sectional design with a sample of 145 women (mean age = 69.9 years), a cumulative ordinal logistic regression model was fitted to MOCA-Col scores, incorporating age and the interaction between the standardized number of children and grandchildren. A higher number of children was significantly associated with lower odds of being in a better cognitive category (β = −0.869, SE = 0.305, *p* = 0.0104, OR = 0.42). Crucially, a significant positive interaction between number of children and number of grandchildren was observed (β = 0.354, SE = 0.093, *p* < 0.001, OR = 1.42), indicating that the negative association between parity and cognition progressively attenuated as the number of grandchildren increased. This evidence supports human life history models and the grandmother hypothesis, suggesting that intergenerational investment may mitigate the cumulative biological costs of reproduction. The findings reflect an evolutionary trade-off between early reproductive effort and later somatic maintenance.

## Introduction

Cognitive decline associated with dementias has shown a sustained increase in prevalence over the years, with Alzheimer’s disease being the most representative condition [1,2]. It has been hypothesized that this increase is related to the rise in life expectancy, supported by access to more hygienic environments and improvements in public health and medicine [3]. As an indirect consequence of this greater longevity, there is also a rise in the incidence of various diseases associated with aging, including late-onset dementias [4,5].

However, this approach does not provide a complete explanation of the issue; as a result, various explanatory perspectives have been proposed. One of these focuses on the study of non-modifiable and modifiable risk factors as precursors to the onset of dementias. The former include characteristics inherent to the individual, such as age, sex, family history, and genetic factors, including the ApoE4 gene. The latter are related to lifestyle factors and psychosocial variables, such as obesity, a sedentary lifestyle, low educational attainment, loneliness, or a lack of social networks in old age, as well as cardiometabolic conditions, such as vascular disease, hypercholesterolemia, diabetes mellitus, and levels of sex hormones in the blood [3,6].

It could thus be expected that non-modifiable risk factors would remain present throughout the population without major differences, due to their very nature. Consequently, one might anticipate a relatively similar trend at the global level, as is the case with age and female sex [7], family history of dementia [4], and genetic factors [8,9]. However, studies have shown that there is no difference between sexes in ages between 65 and 89 years [10], and that this changes by country and continent [11].

Based on the previously described phenomenon, an evolutionary approach could be considered for the study of these pathologies, in which, aside from exploring the mechanisms through which the disease manifests, the ultimate causes of the event are addressed, questioning the reason for its appearance [12]. This approach finds support in the aging processes developed throughout human evolution [13], which involved profound transformations in *Homo sapiens sapiens*, including delayed maturation, slower growth, greater fertility, and the appearance of menopause in women as an indicator of the end of reproductive life [14,15]. In this context, there is currently a recognized link between a longer fertile period and later onset of menopause [16] with delayed cognitive decline, in addition to the positive effects associated with exposure to gonadal steroids, better cognitive performance, and delayed deterioration [17].

By focusing on these issues and on the study of the primate group, it has been observed that they display clear signs and symptoms associated with these pathologies during aging. Among these are, for example, the presence of amyloid plaque deposits and neurocytoskeletal abnormalities, which could be related to the longevity of this group and to the high metabolic demand associated with large brains [4]. Likewise, in both species, cholinergic neurons decrease, and the senile behavioral changes observed in humans and non-human primates are relatively similar [18]. Thus, the extent of these manifestations of structural and cognitive deterioration within this phylogenetic category suggests an evolutionary origin of the disease, which would help explain its presence and development [4,19].

However, it is important to consider that in humans, the cerebral metabolic rate is considerably higher, reaching approximately 20% of total resting energy consumption—twice that observed in apes and up to ten times higher than that of other mammals [14]. This high energy demand correlates with an increase in oxidative stress, which accelerates neuronal aging by affecting the expression of genes directly related to learning, memory, and neuronal survival, thus contributing to the acceleration of pathological aging associated with cognitive decline [20].

In addition, humans are the only primates in which the brain’s metabolic requirement must be maintained over a prolonged period of neurological development [21]. This means that in addition to glucose, which is the primary energy source, it is necessary to draw on additional energy sources to sustain the proper functioning of the nervous system [22]. Nevertheless, it is important to consider that the increase in brain mass has enabled humans to adapt to diverse environmental demands and foster the formation of a cognitive reserve, which acts as a protective factor capable of mitigating the effects of oxidative stress, establishing an inverse relationship between the two processes [23]. Another distinguishing characteristic of humans is the relative size of the brain and development of the neocortex, where late myelination delays the functional maturity of these regions, particularly the frontal lobes [3]. This process is linked to the altricial nature of the human species, in which offspring are more dependent on maternal care [24].

This dependence is associated with an initially low productivity of the offspring and the need for prolonged investment of time and energy in learning and the acquisition of skills and knowledge. In the long term, this pattern contributes to reduced mortality and increased longevity, promoting greater productivity during adulthood [15]. This set of characteristics could have favored the selection of the ApoE allele and late maternal maturity [25].

Regarding genetic characteristics, and specifically the ApoE4 allele, there is currently an explanation for its preservation in the human species based on the concept of antagonistic pleiotropy. This principle suggests that certain genes can have beneficial effects during the early stages of life, whereas in old age, they result in undesirable consequences, a period during which the pressure of natural selection is weaker [8,26,27]. This appears to be the case for the gene associated with the ApoE ε4 allele, which may be responsible for a higher risk of developing Alzheimer’s disease at advanced ages and is recognized as the ancestral form of the gene. The ApoE ε3 and ε2 variants are thought to have appeared later, following the separation of the human lineage from that of chimpanzees and bonobos [3]. Likewise, there is data suggesting that the ApoE ε3 allele is associated with fertility, being more frequent in women who have had children [16]. This could explain the higher population frequency of the ApoE ε3 allele at 79% [28], since it provides a favorable balance that maximizes total reproductive success throughout the lifespan of women, while posing a less severe risk of cognitive problems in old age.

The presence of this gene, and of this allele in particular, is associated with various benefits, among them a reduced risk of spontaneous abortion [29] and greater fertility, mediated by increased progesterone levels. This hormone plays a key role in maintaining the intrauterine environment until the placenta is fully able to meet the needs of the fetus and is also involved in maternal-fetal immunotolerance processes. Additionally, the presence of this allele confers at least three further advantages: cognitive benefits during infant development, protection against infectious diseases, and reproductive advantages [30]. This combination of effects could help explain the persistence and selection of this allele up to the present day.

In contrast, at the molecular level, telomere length and its relationship with the number of children have been studied, and a positive and significant correlation between these variables has been found. In light of these findings, the cooperative breeding strategy in humans has been proposed as a possible explanation, in which a higher number of offspring increases social support—through allomaternal care—thus reducing reproductive costs and the rate of cellular aging. Likewise, a positive influence of Estradiol (E2) has been identified, as its levels increase during pregnancy and act as a protective factor for telomeres against oxidative stress [31]. Continuing with the life history perspective, it is necessary to consider the inherent trade-off between a woman’s reproductive investment and somatic maintenance, as well as the effect this relationship could have on aging. In contexts characterized by resource scarcity and highly restrictive environments, greater reproductive investment could lead to accelerated aging, mediated by increased oxidative stress [32]. In this framework, ApoE4 has been shown to act as a protective factor for reproduction even in hostile environments, which would favor a scenario of antagonistic pleiotropy [33]. Nonetheless, under conditions of resource abundance, this same allele could exert opposite and potentially favorable effects on mothers. In line with this idea, [32] and [34] suggested that oxidative stress could represent the physiological cost of reproductive effort in humans, so that women would assume a greater investment of metabolic energy during reproduction, which would translate into faster aging.

Based on the abovementioned points, it is evident that studying these factors specifically in women is important, given that they represent the population most affected by these changes. In this regard, it has been observed that women who breastfeed for longer periods have a lower risk of developing Alzheimer’s disease [35]. Additionally, evidence has been found that women who have not had children (nulliparous) are exposed throughout their lives to higher levels of estrogen—approximately 22% higher —compared to those who have had offspring [36]. This greater estrogen exposure has been linked to better cognitive performance in old age, attributable to the neuroprotective effect of circulating estrogens [7,37].

In animal studies, it has been observed that estrogens inhibit the formation of beta-amyloid, promote its clearance, reduce neuronal apoptosis, inhibit the hyperphosphorylation of the TAU protein, and decrease both brain oxidative stress and inflammation [35]. Furthermore, they exert direct and indirect effects on various neurotransmitter systems, improve neuronal plasticity, increase cerebral blood flow, and promote the degradation of beta-amyloid precursors [38]. Additionally, reproductive effort has been linked to an increase in oxidative stress experienced by the mother, which, in turn, correlates with the size of the offspring and the duration of the lactation period [39].

Based on the above, the central focus of this research is the analysis of risk factors related to reproductive history, with specific consideration of the number of children [30,40]. From a life history perspective, it is recognized that these trajectories are shaped by environmental conditions and decisions aimed at maximizing the biological advantage associated with passing on genes to future generations, as well as intergenerational support, including contributions to the care of grandchildren, a typically human trait [24]. Likewise, this approach is related to reproductive strategies, which consider the timing within the life cycle when investment in reproduction occurs and the effects that such decisions may have in later stages of life, depending on the evolutionary plasticity of longevity, which responds to previous reproductive decisions [41].

Regarding reproductive history, the number of children a woman has is known to be a risk factor for the development of Alzheimer’s disease. In this respect, it has been reported that women with more than five births are up to 1.7 times more likely to develop the disease. However, these findings cannot be explained solely by variations in estrogen concentrations, as the results differ depending on whether the pregnancies were carried to term, and risk levels vary according to the number of children [42]. Along these lines, [43] found that exposure to both estrogen and progesterone present a protective factor.

Despite this information, other authors have reported divergent findings related to these results [44]. While they agree that estrogen deprivation during the gestational period constitutes a risk factor for the neuropathological development of Alzheimer’s disease, they also point out that an increase in progesterone production could offer benefits by acting as a protector against oxidative stress. These findings suggest the need to further study the variables involved in women’s reproductive history and their relationship with dementia, as well as to consider different sociodemographic conditions, to obtain a more comprehensive explanation of the phenomenon. In this context, the present study does not aim to directly assess the evolutionary mechanisms underlying cognitive decline in old age or to experimentally isolate the specific role of biological factors, such as oxidative stress, sex hormones, or the ApoE allele. Rather, it seeks to empirically analyze the relationship between women’s reproductive history (operationalized through the number of children) and cognitive performance in old age, measured by the MOCA-Col test. From a life history perspective, it is proposed that the number of children constitutes an indirect indicator of accumulated reproductive effort, which could be associated with variations in cognitive functioning in later stages of life. In this sense, the evolutionary framework was used as an interpretive tool to contextualize the observed findings.

## Materials and Methods

### Participants

This paper presents a secondary analysis, corresponding to the follow-up phase (Phase II) of the study *“Progression to mild cognitive impairment and dementia in a group of adults over 50 years of age in Bogotá,”* which initially included 423 participants. The original study was conducted between 2012 and 2017 by the Fundación Universitaria de Ciencias de la Salud (FUCS), the interdisciplinary memory group of Hospital Universitario San José Infantil (HUSJI), and the Faculty of Psychology at Universidad El Bosque. The primary aim of the original study was to estimate the prevalence of cognitive impairment and to identify associated risk factors in adults over 50 years of age residing in Bogotá. For the purposes of the present study, the analysis was restricted to 145 female participants for whom detailed information on reproductive history was collected. Data were accessed for the purposes of this secondary analysis in August, 2025. All data used in this secondary analysis were de-identified. One co-author (A.M.S.) was involved in the original data collection and had access to identifiable participant information at that stage; however, no identifying information was available to or used by the research team during the present analysis.

### Procedure and materials

Data collection was conducted using a comprehensive assessment protocol that included neuropsychological, neurological, and psychiatric clinical evaluations based on well-established procedures, complemented by a structured clinical interview designed to obtain self-reported health history. Cognitive performance was evaluated using the Montreal Cognitive Assessment (MoCA) in its Colombian validated version (MOCA-Col) [45]. For the present analysis, variables related to participants’ reproductive and family history were included, specifically the number of partners, number and sex of children and grandchildren, and age at conception.

The assessment protocols were administered by project researchers, senior undergraduate psychology students, and family medicine residents who had received prior training. All evaluations were conducted at the facilities of the Hospital Universitario San José Infantil. Participation was voluntary; individuals were contacted by telephone and invited to participate in the second phase of the study. Participants attended the assessment accompanied by a family member or friend, and all procedures were preceded by the signing of informed consent, in accordance with ethical principles for research involving human participants and with approval from Fundación Universitaria de Ciencias de la Salud (FUCS) ethics committee (Code: CIDI4909).

MOCA-Col performance was modeled as a function of age, and the interaction between parity (number of children) and number of grandchildren.

Because the numbers of children and grandchildren are conceptually and empirically closely related, both variables were centered and scaled (z-scores) to facilitate interpretation and reduce collinearity issues. Their interaction was included to test whether the association between number of children and cognitive performance varied as a function of the number of grandchildren.

### Statistical analysis

Three modeling approaches were initially evaluated to examine the association between cognitive performance and the explanatory variables: linear regression, Poisson regression, and cumulative ordinal logistic regression. All models included age, number of children, and number of grandchildren as predictors, as well as the interaction between the number of children and the number of grandchildren. Model performance was compared using the Akaike Information Criterion (AIC), and the model with the best relative fit was selected for inference.

Given the ordinal nature of the categorized MOCA outcome, the final model consisted of a cumulative ordinal logistic regression model with a logit link function, assuming proportional odds. The model can be expressed as:

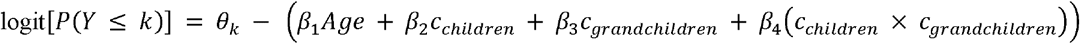

where *Y* represents the ordinal cognitive status, *k* denotes the category thresholds, and θ_*k*_ are the estimated cut points separating the cognitive categories.

For the purposes of the analysis, MOCA scores were categorized into three ordered levels of cognitive status: severe impairment (<18), mild impairment (18–25), and normal cognition (≥26), following established cutoff points for the instrument. This categorization allowed the outcome variable to be analyzed as an ordinal measure of cognitive functioning.

### Model diagnostics

Additional diagnostic analyses were conducted to evaluate the adequacy of the cumulative ordinal model. Potential scale effects were assessed using likelihood ratio tests to determine whether the predictors influenced the scale parameter of the model.

The proportional odds assumption was evaluated using the Brant test, which assesses whether the regression coefficients remain constant across cumulative logits.

### Interaction analysis

To further explore the interaction between the number of children and the number of grandchildren, a Johnson–Neyman procedure was applied. This approach allows identification of the range of values of the moderating variable (number of grandchildren) for which the association between the number of children and cognitive status becomes statistically significant.

### Software

All statistical analyses were performed using the R statistical computing environment [46], employing the package *ordinal* to fit cumulative ordinal regressions [47] and *interactions* to test interactive effects [48]. Figures were made in *ggplot2* [49].

### Data and code availability

All data and code necessary to reproduce the analyses reported in this article are available at the Open Science Framework repository: Sánchez, O. R., Salazar, A. M., & Leongómez, J. D. (2026). *Reproductive history and cognitive aging: Interactive effects of children and grandchildren — Data, Code, and Supplementary Materials*. OSF. https://doi.org/10.17605/OSF.IO/Z53Q2 (CC-BY 4.0).

## Results

Table 1 presents a descriptive summary of the variables used in the study. The analysis was restricted to the 145 female participants in the sample. For continuous variables, the mean, median, and interquartile range (IQR) are reported to characterize their distribution. As detailed, the cohort had a mean age of 69.9 years, with a mean of 3.1 children and 4.2 grandchildren, and an average cognitive performance, measured by MOCA-Col, of 21.3 points.

**Table 1.** Descriptive characteristics of the study population.

| Variable | Mean (SD) | Median (IQR) |
| --- | --- | --- |
| Age (years) | 69.9 ± 6.8 | 69 (9) |
| MOCACol (Score) | 21.3 ± 4.9 | 22 (7) |
| Number of children | 3.1 ± 2.9 | 3 (2.5) |
| Number of grandchildren | 4.2 ± 5.5 | 3 (5.25) |

Before presenting the model that best fits the data, a correlation analysis was performed between the number of children and the number of grandchildren, which resulted in a high correlation (0.80). To avoid collinearity, the decision was made to center and scale the variables, and then to find the best fit for the model.

Three modeling approaches were evaluated to examine the association between MOCA score and demographic and family-related variables: (1) linear regression, (2) Poisson regression, and (3) cumulative ordinal logistic regression.

All models included age, number of children, and number of grandchildren (centered variables), as well as the interaction between the number of children and grandchildren.

Model performance was compared using the Akaike Information Criterion (AIC) (Table 2), and the model with the best relative fit was selected.

**Table 2.** Model comparison using Akaike Information Criterion.

| Model | AIC | ΔAIC |
| --- | --- | --- |
| Cumulative ordinal model | 243.10 | 0.0000 |
| Linear regression | 828.87 | 585.77 |
| Poisson Regression | 841.82 | 598.72 |

**Note:** AIC = Akaike Information Criterion. ΔAIC represents the difference between the AIC of each model and the lowest AIC observed among the candidate models. Lower AIC values indicate better model fit.

The cumulative ordinal model showed the lowest AIC, indicating a substantially better fit compared with the linear and Poisson regression models.

A cumulative ordinal regression model (Table 3) was fitted to evaluate the association between age, number of children, and number of grandchildren with cognitive status categorized according to MOCA scores (severe impairment, mild impairment, and normal cognition). The proportional odds model with logit link showed good convergence after five iterations and included 142 observations (AIC = 243.10).

**Table 3.** Cumulative ordinal logistic regression model for cognitive impairment (MOCA-Col)

| Section | Variable | Beta | SE | z | p | OR | LI | LS |
| --- | --- | --- | --- | --- | --- | --- | --- | --- |
| Predictors | Age | -0.077 | 0.028 | -2.76 | 0.006 | 0.92 | 0.88 | 0.98 |
| Predictors | Number of children | -0.869 | 0.305 | -2.85 | 0.004 | 0.42 | 0.23 | 0.76 |
| Predictors | Number of grandchildren | -0.810 | 0.333 | -2.43 | 0.015 | 0.45 | 0.23 | 0.85 |
| Predictors | Number of children ×<br>Number of grandchildren | 0.354 | 0.093 | 3.82 | <0.001 | 1.42 | 1.19 | 1.71 |
| Thresholds | Severe \| Mild | -6.848 | 2.004 | -3.42 | <0.001 |  |  |  |
| Thresholds | Mild \| Normal | -3.385 | 1.925 | -1.76 | 0.079 |  |  |  |

**Note:** OR = Odds ratio. Threshold parameters represent the estimated cut-points separating adjacent categories of the ordinal outcome (MOCA cognitive status).

Age was significantly associated with cognitive status. Each additional year of age was associated with lower odds of being in a better cognitive category (β = −0.077, SE = 0.028, *p* = 0.006), corresponding to an odds ratio (OR) of 0.92.

Similarly, a higher number of children was associated with poorer cognitive outcomes (β = −0.869, SE = 0.305, *p* = 0.004), corresponding to an OR of 0.42, indicating lower odds of being in a higher cognitive category. A comparable association was observed for the number of grandchildren (β = −0.810, SE = 0.333, *p* = 0.015; OR = 0.45).

Importantly, a statistically significant interaction between number of children and number of grandchildren was observed (β = 0.354, SE = 0.093, *p* < 0.001), corresponding to an OR of 1.42. This suggests that the association between fertility history and cognitive status depends on the combined effect of offspring across generations.

The estimated threshold parameters separating severe from mild cognitive impairment (β = −6.848) and mild impairment from normal cognition (β = −3.385) were consistent with the ordinal structure of the outcome. The model is summarized in Fig 1.

**Fig 1.**
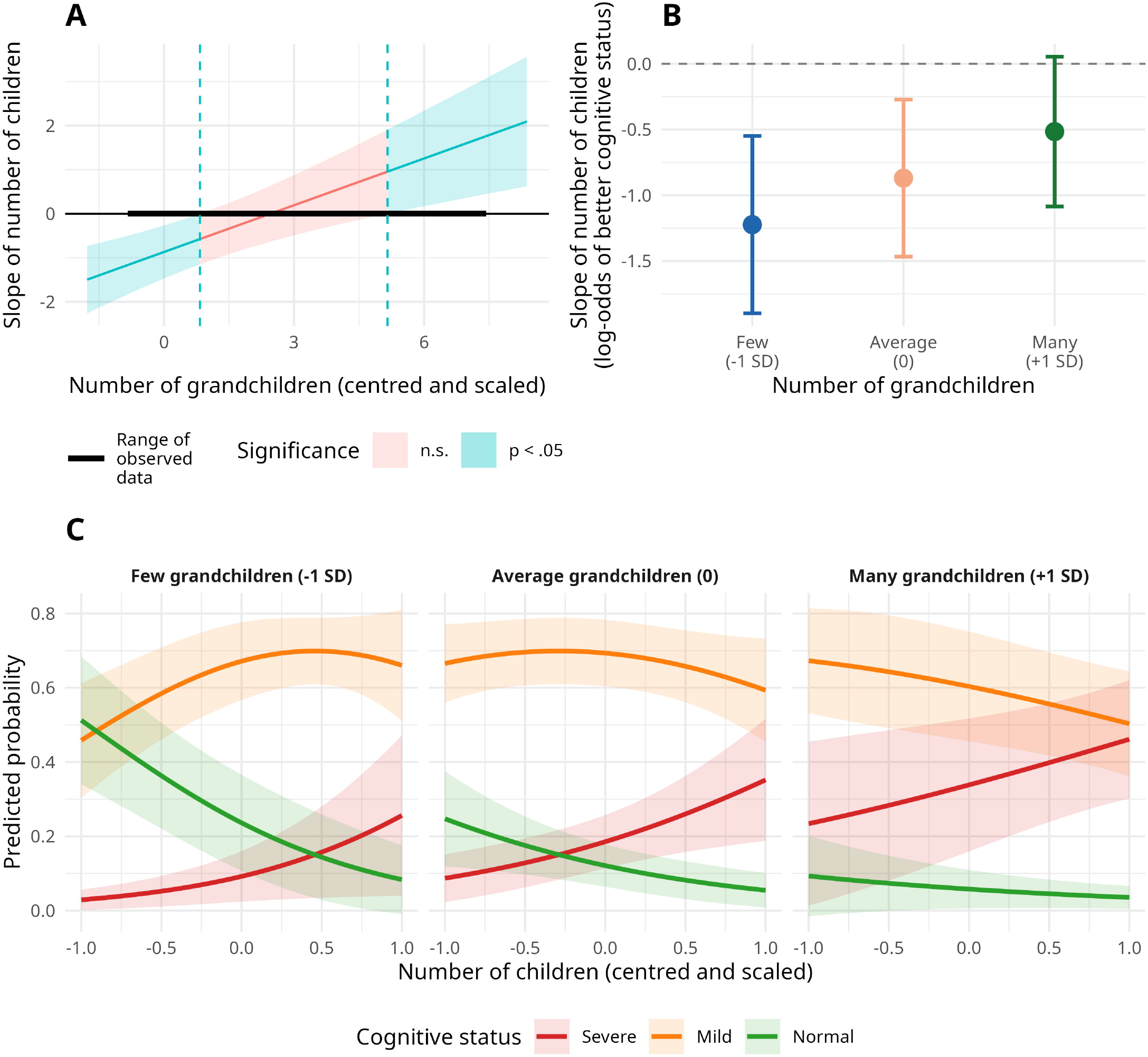
Interaction between number of children and number of grandchildren on cognitive status. (A) Johnson–Neyman plot showing the range of number of grandchildren (centered and scaled) for which the association with number of children is statistically significant (blue shading, p < 0.05) or not (red shading, p > 0.05). Dashed vertical lines mark the significance boundaries. (B) Simple slopes of the effect of number of children on the log-odds of better cognitive status at three levels of number of grandchildren. Points represent slope estimates; error bars represent 95% confidence intervals. The dashed horizontal line indicates a slope of zero. (C) Predicted probabilities of each cognitive status category (Severe, Mild, Normal) as a function of number of children (centered and scaled), separately for low (-1 SD), average (0), and high (+1 SD) levels of number of grandchildren.

At low levels of grandchildren (−1 SD), the effect of number of children is most pronounced (Fig 1C): the probability of normal cognition decreases sharply as the number of children increases, while the probability of severe impairment rises markedly, reflecting the most unfavorable scenario. At average levels of grandchildren (0), the same directional pattern is observed but is considerably attenuated, with mild impairment remaining the dominant category across the full range of children. At high levels of grandchildren (+1 SD), the probability of severe impairment continues to increase with number of children, but at a substantially slower rate, consistent with the moderating role of grandchildren identified in the regression model. Notably, the probability of normal cognition is low across all levels of grandchildren in this sample, which likely reflects the confounding influence of age — older women are both more likely to have more grandchildren and less likely to show normal cognitive performance.

Taken together, these results indicate that the reproductive cost associated with a higher number of children is expressed primarily through the transition between severe and mild impairment categories, and that this cost is progressively attenuated in the presence of more grandchildren.

Additional diagnostic analyses were performed to assess potential scale effects in the cumulative ordinal model. Likelihood ratio tests indicated no evidence that age, number of children, number of grandchildren, or their interaction influenced the scale parameter of the model (all *p* > 0.10), supporting the adequacy of the proportional odds specification.

The proportional odds assumption was evaluated using the Brant test, which indicated no evidence of violation of the proportional odds assumption (global test *p* > 0.05).

The Johnson–Neyman analysis (Fig 1A) was used to further examine the interaction between the number of children and the number of grandchildren on cognitive status. As shown in Figure 1, the effect of the number of children on cognitive status varied depending on the number of grandchildren. Specifically, the effect of number of children on cognitive status was statistically significant only at relatively low values of number of grandchildren. Beyond the upper boundary, the association attenuated and ceased to be significant. This pattern indicates that the relationship between fertility history and cognitive status is conditional on the number of grandchildren, supporting the interaction observed in the cumulative ordinal regression model. The interval where the effect is not significant is quite wide, suggesting that the moderating effect of grandchildren is gradual.

## Discussion

The results showed that the number of children was associated with lower odds of being in a better cognitive category, which is related to the findings of [36], who reported that nulliparous women are exposed to higher concentrations of estrogen throughout their lives, representing a protective factor [7,37]. This finding aligns with the reproductive cost hypothesis, which suggests that reproductive investment may contribute to long-term health consequences in old age, though our cross-sectional design limits causal inference.

However, the most significant finding is the countervailing effect observed in the interaction with the number of grandchildren: a higher number of grandchildren is associated with a protective effect on cognitive performance, even among women with high parity. This finding suggests a potential mechanism where accumulated reproductive costs may be partially offset by benefits associated with intergenerational investment, an interpretation consistent with human life history models and the grandmother hypothesis.

Our species has a distinctive characteristic compared to other primates: postmenopausal life [50–52], a critical component of the human life history strategy. The adaptive significance of grandparents lies not only in the transmission of knowledge and skills but also in providing resources to children and grandchildren, which can affect offspring survival [15,53]. This feature appears to be related to the altricial condition of human infants [54], which demands intense dedication from mothers, reducing the time available for weaned children, who receive direct support from their grandmothers [55]. In this sense, postmenopausal life could represent an adaptive advantage associated with the slowing of aging [56], increasing the biological efficacy of both mothers and grandmothers, with indirect effects on the fertility of younger generations [50]. The rate of aging, therefore, could be linked to the contribution of adults to intergenerational juvenile survival (Kaplan & Robson, 2009), with evolutionary implications for the accumulation of deleterious mutations [58].

These considerations make sense when considering the potential protective factors associated with being a grandmother or grandfather. While caring for grandchildren can involve costs related to the time invested, physical effort, and health impacts in low-resource settings [59], especially when the children are young—which may lead to loss of sleep and greater exposure to infections—long-term benefits have also been described.

While some studies report negative health effects for caregiving grandparents, such as higher rates of depression and hypertension [60], these findings highlight the importance of balancing caregiving demands with available resources, which is crucial for understanding how intergenerational investment can mitigate reproductive costs, as suggested by our results. However, a large-scale longitudinal study showed that although a decline was initially observed, the trend changed over time, suggesting a positive long-term effect of caregiving [61].

In a study by [62], when comparing caregiving grandmothers in rural and urban areas, the main health problems reported were arthritis (over 50%), followed by diabetes (19%) and poor circulation (18%). It is noteworthy that Alzheimer’s ranked last, with prevalence rates of 1.1% in rural areas and 0.3% in urban areas, below those for more than ten additional health problems.

Protective factors associated with caregiving activities that promote mental health and well-being have been identified [63] and can be grouped into three categories: personal assets (self-concept and coping skills), social assets (access to health services, religiosity, and support networks), and environmental assets (healthy physical environments, leisure, and financial availability). For grandchild care to have positive effects on grandparents’ health, it is essential that there is a favorable balance between caregiving demands and available resources [61], which is further strengthened by positive personality traits and a valued perception of the caregiver role [64].

It is also relevant to consider the intensity of care, understood as the amount of time dedicated to caring for grandchildren, which is associated with health-related quality of life [65], within the framework of the Family Stress, Resilience, and Adaptation Model [66]. In Europe, an increase in life expectancy has been identified, associated with an increase in the number of grandparent caregivers [67]. Thus, 52% of grandparents aged > 50 years were identified as supplementary caregivers of grandchildren under 16 years old [68]. The intensity of care varies significantly between countries, being higher in Mediterranean regions than in Northern Europe [69].

Determinants of regular caregiving include being female, marital status (married), availability of financial resources, and good health [70]. When demands exceed abilities, caregiving can become a stressor with negative consequences [69]; however, greater benefits have been reported in complementary caregiving contexts in various European countries [71].

Noriega et al. [65] found that the number of grandchildren was associated with an increase in social interaction, although not with greater subjective satisfaction. This suggests that the relevant variable is not merely the number of grandchildren, but rather the quality of the relationships established. Nevertheless, having more grandchildren could increase opportunities to form close and meaningful bonds.

Additionally, the strength of character in grandparents has been identified as a protective factor that positively influences the quality of life related to physical and mental health during the care of grandchildren [65].

Specific studies have examined the impact of caring for grandchildren on cognitive functioning [72]. In China, where intergenerational responsibility is a core value, 58% of grandparents participate in caring for their grandchildren [73]. These family interactions can be considered forms of cognitive stimulation associated with maintaining cognitive health in old age [74], as they promote neurogenesis, intellectual engagement, and delay cognitive decline. This positive perspective on family caregiving is consistent with role enhancement theory, which suggests that social participation generates emotional satisfaction, improves subjective well-being, and, as a result, may help delay cognitive decline [75].

Our results support the hypothesis that reproductive history entails cumulative biological costs for cognitive health in old age, but that these costs may be partially modulated by the adaptive benefits associated with intergenerational investment. This pattern is consistent with the predictions of human life history theory and suggests that cognitive decline in old age could be understood, at least in part, as the result of evolutionary trade-offs between early reproduction and late-life somatic maintenance.

### Study limitations

This study has some limitations that should be considered when interpreting the results. First, the cross-sectional design prevents us from establishing causal relationships between women’s reproductive history and cognitive performance in old age; therefore, the observed associations do not allow us to infer temporal directionality or completely rule out the possibility of reverse causality or unmeasured confounding variables. Second, although the MOCA-Col is a widely validated screening instrument, it does not allow for the differentiation of specific neuropsychological profiles or the establishment of clinical diagnoses of dementia; therefore, the results should be interpreted as indicators of overall cognitive functioning and not as the presence or absence of neurodegenerative disease. Third, women’s reproductive history was assessed using self-report, which may be subject to recall bias, particularly in older populations. Fourth, no *a priori* power analysis was conducted, and no equivalence tests were performed, as this study is based on a secondary analysis of previously collected data with a fixed sample size. Accordingly, the findings should be regarded as exploratory rather than confirmatory, and non-significant results should not be interpreted as evidence in favor of the null hypothesis. Finally, the results are based on a specific sample from Bogotá, Colombia, and may not be generalizable to populations with different cultural, economic, and family contexts. Intergenerational caregiving practices, for example, vary considerably across regions and cultures, suggesting that the observed protective effects of grandchildren might be amplified or diminished in other settings. Future longitudinal studies that integrate biological markers, contextual variables, and more detailed neuropsychological measures will allow for a more accurate assessment of the underlying mechanisms and directionality of the observed associations.

## Supporting information

https://osf.io/z53q2/overview?view_only=1cd45ce2171d4accb2309341b4c4b8e2

## Acknowledgments

We express our sincere gratitude to the Fundación Universitaria de Ciencias de la Salud (FUCS) and the Interdisciplinary Memory Group of the Hospital Infantil Universitario San José (HIUSJ) for their valuable support in the development of this work. We especially acknowledge the collaboration and commitment of Dr. Olga Lucía Pedraza Linares, whose guidance was fundamental throughout the research process.

