## Supplementary material for "Reproductive history and cognitive aging: Interactive effects of children and grandchildren in a life history framework": https://osf.io/z53q2/overview?view_only=1cd45ce2171d4accb2309341b4c4b8e2

Supplementary Materials: Code and Analyses

Oscar R. Sánchez[1*]    Ana María Salazar[2]    Juan David Leongómez[1]

9 April, 2026

[1] CODEC: Cognitive and Behavioural Sciences Research Group, Faculty of Psychology, Universidad El Bosque, Bogotá 110121, Colombia

[2] Health, Sports, and Clinical Psychology, Faculty of Psychology, Universidad El Bosque, Bogotá 110121, Colombia

**8 Manuscript figure** 26

**9 Session information** 30

### 1 Description

This document contains all code and step-by-step explanations for the analyses, figures, and tables reported in:

> Sánchez, O. R., Salazar, A. M., & Leongómez, J. D. (under review). *Reproductive history and cognitive aging: Interactive effects of children and grandchildren in a life history framework.*

The document is organised as follows. After loading packages and data, we fit three candidate models: linear regression, Poisson regression, and cumulative ordinal logistic regression, in that order. We then compare them using the Akaike Information Criterion (AIC) and justify the selection of the ordinal model as the primary model reported in the manuscript. Finally, we present the full diagnostic and inferential results for that model, including the interaction analysis via Johnson–Neyman procedure and simple slopes.

---

### 2 Preliminaries

#### 2.1 Load packages

```
library(MASS) # polr (for Brant test)
library(tidyverse) # data wrangling and figures
library(broom) # tidy model output
library(flextable) # publication tables
library(emmeans) # estimated marginal means and contrasts
library(ordinal) # cumulative link models
library(interactions) # interaction analysis (Johnson-Neyman)
```

```
library(AER) # dispersion test
library(car) # VIF
library(brant) # proportional odds test
```

#### 2.2 Load and prepare data

```
data_women <- suppressMessages(
  read_delim("Data/Database.csv", delim = NULL)
) |>
  dplyr::filter(Sex == "Female") |>
  mutate(
    `Number of children`     = as.numeric(as.character(`Number of children`)),
    c_children               = as.numeric(scale(`Number of children`)),
    c_grandchildren          = as.numeric(scale(`Number of grandchildren`))
  )
```

The predictor variables (number of children and number of grandchildren) were centred and scaled (z-scores) prior to modelling. This reduces collinearity between the main effects and their interaction term, and makes coefficients directly interpretable as the effect of a one-standard-deviation change in each variable.

#### 2.3 Descriptive statistics

```
desc <- data_women |>
  summarise(
    across(
      c(Age, MOCACol, `Number of children`, `Number of grandchildren`),
      list(
        mean   = \(x) mean(x, na.rm = TRUE),
        sd     = \(x) sd(x, na.rm = TRUE),
        median = \(x) median(x, na.rm = TRUE),
        IQR    = \(x) IQR(x, na.rm = TRUE)
      )
    )
  )

tibble(
  Variable = c(
    "Age (years)", "MOCA-Col (score)",
    "Number of children", "Number of grandchildren"
```

```
  ),
  `Mean (SD)` = c(
    paste0(
      round(desc$Age_mean, 1),
      " (", round(desc$Age_sd, 1), ")"
    ),
    paste0(
      round(desc$MOCACol_mean, 1),
      " (", round(desc$MOCACol_sd, 1), ")"
    ),
    paste0(
      round(desc$`Number of children_mean`, 1),
      " (", round(desc$`Number of children_sd`, 1), ")"
    ),
    paste0(
      round(desc$`Number of grandchildren_mean`, 1),
      " (", round(desc$`Number of grandchildren_sd`, 1), ")"
    )
  ),
  `Median (IQR)` = c(
    paste0(
      round(desc$Age_median, 1),
      " (", round(desc$Age_IQR, 1), ")"
    ),
    paste0(
      round(desc$MOCACol_median, 1),
      " (", round(desc$MOCACol_IQR, 1), ")"
    ),
    paste0(
      round(desc$`Number of children_median`, 1),
      " (", round(desc$`Number of children_IQR`, 1), ")"
    ),
    paste0(
      round(desc$`Number of grandchildren_median`, 1),
      " (", round(desc$`Number of grandchildren_IQR`, 1), ")"
    )
  )
) |>
  flextable() |>
```

```
  autofit() |>
  theme_booktabs()
```

**Table S1.** Descriptive statistics of the study sample (N = 145 women).

| Variable | Mean (SD) | Median (IQR) |
|---|---|---|
| Age (years) | 69.9 (6.8) | 69 (9) |
| MOCA-Col (score) | 21.3 (4.9) | 22 (7) |
| Number of children | 3.1 (2.9) | 3 (2.5) |
| Number of grandchildren | 4.2 (5.5) | 3 (5.2) |

---

### 3 Model 1: Linear regression

As a baseline, we fitted an ordinary least-squares linear regression of MOCA-Col scores on age, centred number of children, centred number of grandchildren, and their interaction.

#### 3.1 Fit

```
model_lm <- lm(
  MOCACol ~ Age + c_children * c_grandchildren,
  data = data_women
)

summary(model_lm)
```

```

Call:
lm(formula = MOCACol ~ Age + c_children * c_grandchildren, data = data_women)

Residuals:
     Min       1Q   Median       3Q      Max
-13.3510  -2.4229   0.3582   3.2412  10.3313

Coefficients:
                           Estimate Std. Error t value Pr(>|t|)
(Intercept)                34.61221    4.04713   8.552 2.14e-14 ***
Age                        -0.19767    0.05748  -3.439 0.000774 ***
```

```
c_children                   -1.39295    0.62907  -2.214 0.028462 *
c_grandchildren              -1.34095    0.70135  -1.912 0.057968 .
c_children:c_grandchildren    0.60594    0.18590   3.260 0.001407 **
---
Signif. codes:  0 '***' 0.001 '**' 0.01 '*' 0.05 '.' 0.1 ' ' 1

Residual standard error: 4.372 on 137 degrees of freedom
  (3 observations deleted due to missingness)
Multiple R-squared:  0.2162,	Adjusted R-squared:  0.1933
F-statistic: 9.447 on 4 and 137 DF,  p-value: 8.952e-07
```

##### 3.2 Coefficients

```r
tidy(model_lm) |>
  mutate(
    estimate = round(estimate, 3),
    std.error = round(std.error, 3),
    statistic = round(statistic, 2),
    p.value = ifelse(p.value < .001, "< .001",
      sub("^0", "", sprintf("%.3f", p.value))
    )
  ) |>
  rename(
    Predictor = term, B = estimate, SE = std.error,
    t = statistic, p = p.value
  ) |>
  flextable() |>
  autofit() |>
  theme_booktabs()
```

**Table S2.** Linear regression model for MOCA-Col scores.

| Predictor | B | SE | t | p |
|---|---|---|---|---|
| (Intercept) | 34.612 | 4.047 | 8.55 | < .001 |
| Age | -0.198 | 0.057 | -3.44 | < .001 |
| c_children | -1.393 | 0.629 | -2.21 | .028 |
| c_grandchildren | -1.341 | 0.701 | -1.91 | .058 |
| c_children:c_grandchildren | 0.606 | 0.186 | 3.26 | .001 |

| Predictor | B | SE | t p |
|---|---|---|---|

##### 3.3 Residual diagnostics

```
par(mfrow = c(2, 2))
plot(model_lm)
par(mfrow = c(1, 1))
```

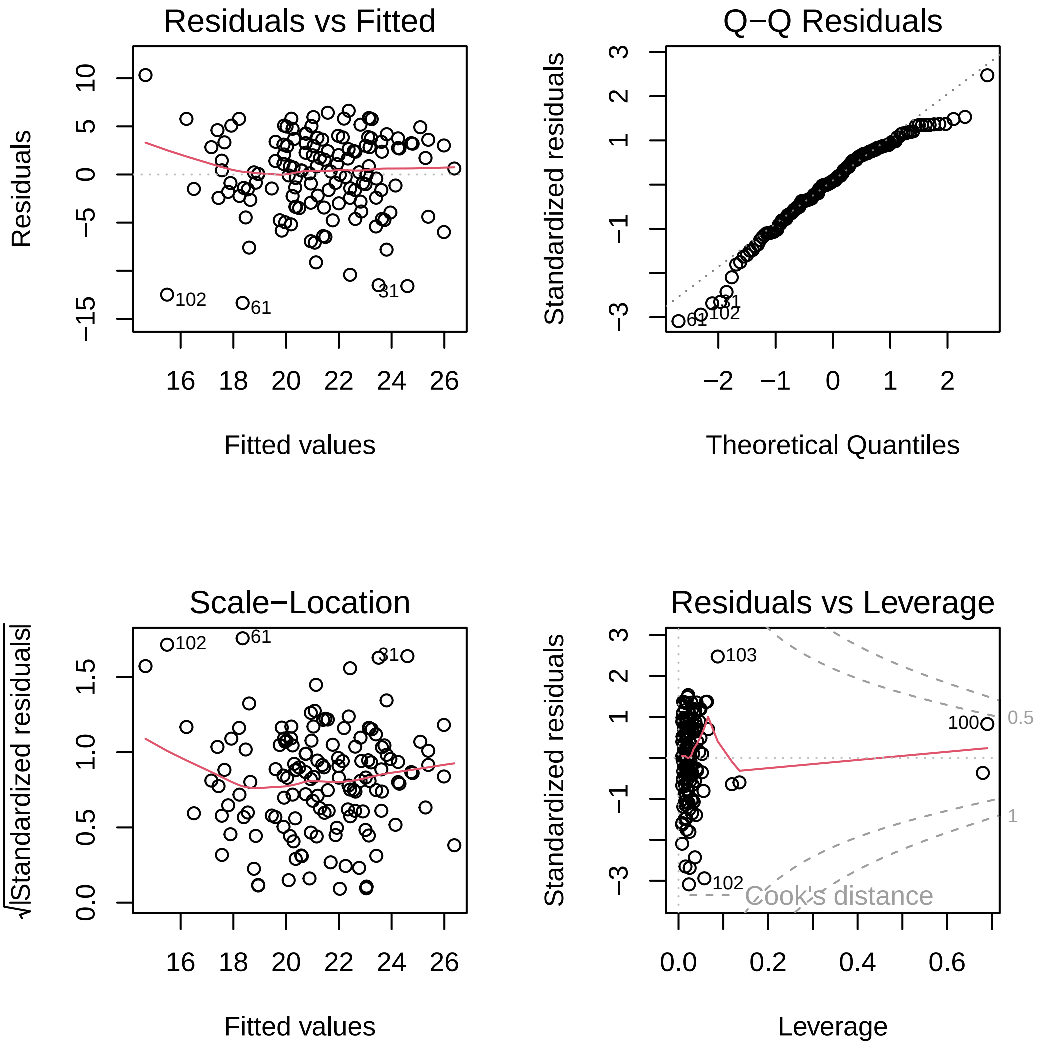

**Figure S1.** Residual diagnostic plots for the linear regression model.

##### 3.4 Predicted values

```r
newdat_lm <- expand.grid(
  c_children      = seq(-3, 3, length.out = 100),
  c_grandchildren = c(-1, 0, 1),
  Age             = mean(data_women$Age, na.rm = TRUE)
)

newdat_lm$MOCA_pred <- predict(model_lm, newdata = newdat_lm)

newdat_lm |>
  mutate(
    Grandchildren = factor(c_grandchildren,
      levels = c(-1, 0, 1),
      labels = c(
        "Few grandchildren (-1 SD)",
        "Average grandchildren (0)",
        "Many grandchildren (+1 SD)"
      )
    )
  ) |>
  ggplot(aes(x = c_children, y = MOCA_pred, colour = Grandchildren)) +
  geom_line(linewidth = 1.1) +
  scale_colour_manual(values = c("#2166ac", "#f4a582", "#1a7837")) +
  labs(
    x      = "Number of children (centred and scaled)",
    y      = "Predicted MOCA-Col score",
    colour = NULL
  ) +
  theme_minimal(base_size = 12) +
  theme(legend.position = "bottom")
```

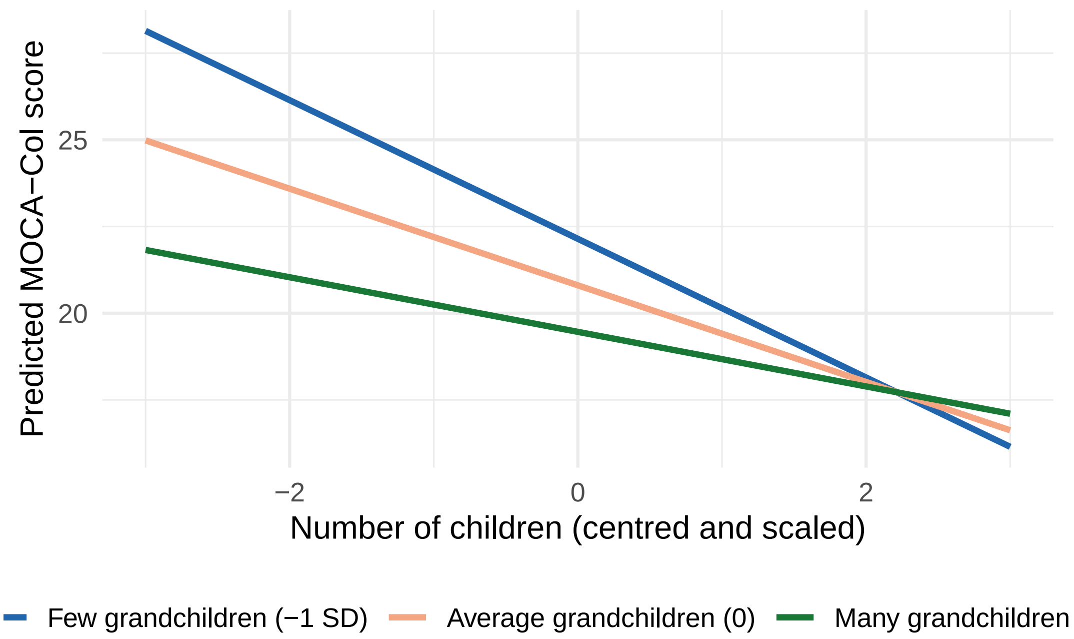

**Figure S2.** Predicted MOCA-Col scores from the linear model as a function of number of children (centred), at three levels of number of grandchildren.

---

### 4 Model 2: Poisson regression

Because MOCA-Col scores are non-negative counts, we also fitted a Poisson generalised linear model with the same predictor structure.

#### 4.1 Fit

```
model_pois <- glm(
  MOCACol ~ Age + c_children * c_grandchildren,
  family = poisson,
  data   = data_women
)

summary(model_pois)
```

```
Call:
glm(formula = MOCACol ~ Age + c_children * c_grandchildren, family = poisson,
    data = data_women)
```

```
Coefficients:
                             Estimate Std. Error z value Pr(>|z|)
(Intercept)                  3.680105   0.199570  18.440  < 2e-16 ***
Age                         -0.009316   0.002850  -3.269  0.00108 **
c_children                  -0.066238   0.031334  -2.114  0.03452 *
c_grandchildren             -0.064976   0.035285  -1.841  0.06556 .
c_children:c_grandchildren  0.028923   0.009193   3.146  0.00165 **
---
Signif. codes:  0 '***' 0.001 '**' 0.01 '*' 0.05 '.' 0.1 ' ' 1

(Dispersion parameter for poisson family taken to be 1)

    Null deviance: 174.30  on 141  degrees of freedom
Residual deviance: 140.07  on 137  degrees of freedom
  (3 observations deleted due to missingness)
AIC: 841.82

Number of Fisher Scoring iterations: 4
```

##### 4.2 Coefficients

```
tidy(model_pois) |>
  mutate(
    estimate = round(estimate, 4),
    std.error = round(std.error, 4),
    statistic = round(statistic, 2),
    p.value = ifelse(p.value < .001, "< .001",
      sub("^0", "", sprintf("%.3f", p.value))
    )
  ) |>
  rename(
    Predictor = term, B = estimate, SE = std.error,
    z = statistic, p = p.value
  ) |>
  flextable() |>
  autofit() |>
  theme_booktabs()
```

**Table S3.** Poisson regression model for MOCA-Col scores.

| Predictor | B | SE | z p |
|---|---|---|---|
| (Intercept) | 3.6801 | 0.1996 | 18.44 < .001 |
| Age | -0.0093 | 0.0028 | -3.27 .001 |
| c_children | -0.0662 | 0.0313 | -2.11 .035 |
| c_grandchildren | -0.0650 | 0.0353 | -1.84 .066 |
| c_children:c_grandchildren | 0.0289 | 0.0092 | 3.15 .002 |

##### 4.3 Dispersion check

A key assumption of Poisson regression is equidispersion (variance = mean). We tested this formally.

```r
# Ratio of residual deviance to degrees of freedom (should be ~1)
cat(
  "Deviance / df:",
  round(deviance(model_pois) / df.residual(model_pois), 3), "\n"
)
```

```
Deviance / df: 1.022
```

```r
# Formal dispersion test
dispersiontest(model_pois)
```

```

	Overdispersion test

data:  model_pois
z = -0.77043, p-value = 0.7795
alternative hypothesis: true dispersion is greater than 1
sample estimates:
dispersion
 0.8934963
```

There is no evidence of overdispersion ($\underline{p}$ = .78); the model is slightly underdispersed (dispersion 0.89), which remains acceptable and does not invalidate the Poisson specification.

##### 4.4 Collinearity (VIF)

```
vif(model_pois)
```

```
                   Age                   c_children 
              1.033486                     2.852256 
        c_grandchildren c_children:c_grandchildren 
              3.739175                     2.586417 
```

All VIF values are below 4 after centring, indicating no problematic collinearity, even given the moderate correlation between number of children and number of grandchildren ($r$ = .80).

##### 4.5 Predicted values

```
newdat_pois <- expand.grid(
  c_children      = seq(-3, 3, length.out = 100),
  c_grandchildren = c(-1, 0, 1),
  Age             = mean(data_women$Age, na.rm = TRUE)
)

newdat_pois$MOCA_pred <- predict(model_pois,
  newdata = newdat_pois, type = "response"
)

newdat_pois |>
  mutate(
    Grandchildren = factor(c_grandchildren,
      levels = c(-1, 0, 1),
      labels = c(
        "Few grandchildren (-1 SD)",
        "Average grandchildren (0)",
        "Many grandchildren (+1 SD)"
      )
    )
  ) |>
  ggplot(aes(x = c_children, y = MOCA_pred, colour = Grandchildren)) +
  geom_line(linewidth = 1.1) +
  scale_colour_manual(values = c("#2166ac", "#f4a582", "#1a7837")) +
  labs(
    x      = "Number of children (centred and scaled)",
    y      = "Predicted MOCA-Col score",
    colour = NULL
```

```
  ) +
  theme_minimal(base_size = 12) +
  theme(legend.position = "bottom")
```

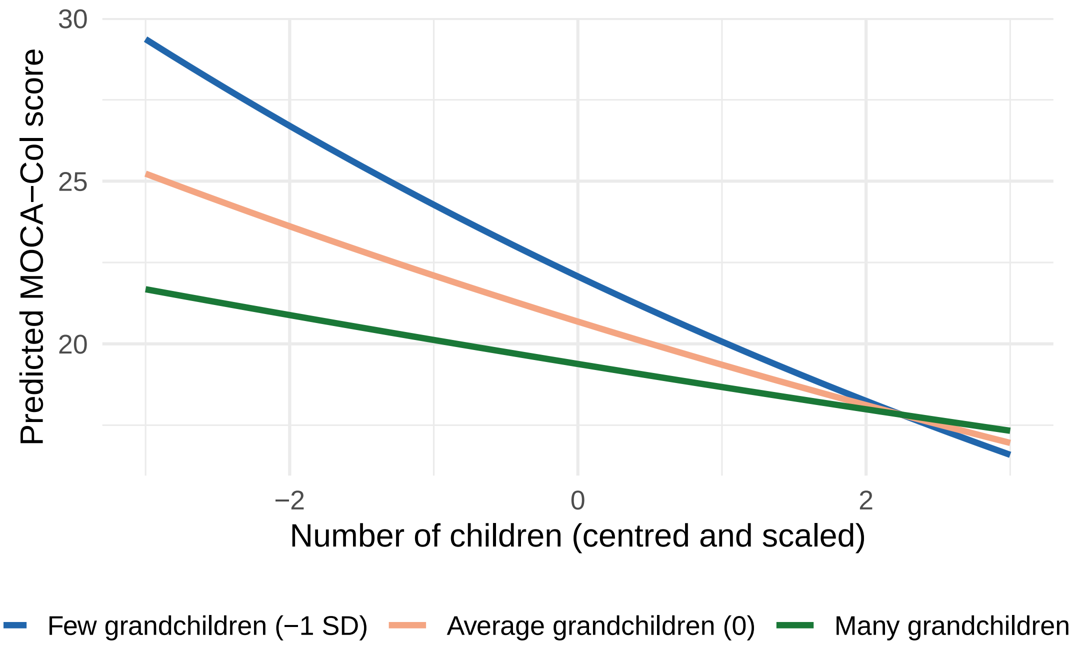

**Figure S3.** Predicted MOCA-Col scores from the Poisson model as a function of number of children (centred), at three levels of number of grandchildren.

---

#### 5 Model 3: Cumulative ordinal logistic regression

Given the ordinal nature of cognitive status, we categorised MOCA-Col scores into three ordered levels: severe impairment (<18), mild impairment (18–25), and normal cognition ( 26), and fitted a cumulative link model (CLM) with a logit link and proportional-odds structure.

##### 5.1 Outcome categorisation

```
data_women <- data_women |>
  mutate(
    MOCA_cat = cut(MOCACol,
      breaks         = c(-Inf, 17, 25, 30),
      labels         = c("Severe", "Mild", "Normal"),
      ordered_result = TRUE
    )
```

```r
  )

table(data_women$MOCA_cat)
```

```
Severe   Mild Normal 
    30     86     29 
```

##### 5.2 Fit

```r
model_clm <- clm(
  MOCA_cat ~ Age + c_children * c_grandchildren,
  data = data_women
)

summary(model_clm)
```

```
formula: MOCA_cat ~ Age + c_children * c_grandchildren
data:    data_women

 link  threshold nobs logLik  AIC    niter max.grad cond.H 
 logit flexible  142  -115.55 243.10 6(0)  2.59e-11 1.3e+06

Coefficients:
                           Estimate Std. Error z value Pr(>|z|)    
Age                        -0.07701    0.02788  -2.762 0.005737 ** 
c_children                 -0.86947    0.30468  -2.854 0.004321 ** 
c_grandchildren            -0.80980    0.33259  -2.435 0.014900 *  
c_children:c_grandchildren  0.35396    0.09270   3.818 0.000134 ***
---
Signif. codes:  0 '***' 0.001 '**' 0.01 '*' 0.05 '.' 0.1 ' ' 1

Threshold coefficients:
            Estimate Std. Error z value
Severe|Mild   -6.848      2.004  -3.417
Mild|Normal   -3.385      1.925  -1.759
(3 observations deleted due to missingness)
```

##### 5.3 Coefficients

```
coefs <- summary(model_clm)$coefficients |>
  as.data.frame() |>
  rownames_to_column("term") |>
  rename(
    estimate = Estimate,
    std.error = `Std. Error`,
    statistic = `z value`,
    p.value = `Pr(>|z|)`
  )

coefs |>
  mutate(
    Section = ifelse(grepl("\\|", term),
      "Thresholds", "Predictors"
    ),
    OR = ifelse(grepl("\\|", term),
      NA_real_, round(exp(estimate), 3)
    ),
    LI = ifelse(grepl("\\|", term),
      NA_real_, round(exp(estimate - 1.96 * std.error), 3)
    ),
    LS = ifelse(grepl("\\|", term),
      NA_real_, round(exp(estimate + 1.96 * std.error), 3)
    ),
    estimate = round(estimate, 3),
    std.error = round(std.error, 3),
    statistic = round(statistic, 2),
    p.value = ifelse(p.value < .001, "< .001",
      sub("^0", "", sprintf("%.3f", p.value))
    )
  ) |>
  arrange(Section) |>  # "Predictors" sorts before "Thresholds" alphabetically
  dplyr::select(
    Section, term, estimate, std.error,
    statistic, p.value, OR, LI, LS
  ) |>
  rename(
    ` ` = Section, Predictor = term, B = estimate, SE = std.error,
```

```
    z = statistic, p = p.value
  ) |>
  flextable() |>
  merge_v(j = " ") |>
  theme_booktabs() |>
  fontsize(size = 9, part = "all") |>
  fit_to_width(max_width = 6.5) |>
  add_footer_lines(
    "Note. OR = odds ratio; LI/LS = 95% CI bounds. Threshold parameters represent
    cut-points separating adjacent cognitive categories."
  )
```

**Table S4.** Cumulative ordinal logistic regression model for cognitive status (MOCA-Col). OR = odds ratio; LI/LS = lower/upper bound of the 95% confidence interval.

| | Predictor | B | SE | z | p | OR | LI | LS |
|---|---|---|---|---|---|---|---|---|
| | Age | -0.077 | 0.028 | -2.76 | .006 | 0.926 | 0.877 | 0.978 |
| Predictors | c_children | -0.869 | 0.305 | -2.85 | .004 | 0.419 | 0.231 | 0.762 |
| | c_grandchildren | -0.810 | 0.333 | -2.43 | .015 | 0.445 | 0.232 | 0.854 |
| | c_children:c_grandchildren | 0.354 | 0.093 | 3.82 | < .001 | 1.425 | 1.188 | 1.709 |
| Thresholds | Severe\|Mild | -6.848 | 2.004 | -3.42 | < .001 | | | |
| | Mild\|Normal | -3.385 | 1.925 | -1.76 | .079 | | | |

Note. OR = odds ratio; LI/LS = 95% CI bounds. Threshold parameters represent cut-points separating adjacent cognitive categories.

#### 5.4 Predicted probabilities

```
newdat_clm <- expand.grid(
  c_children      = seq(-3, 3, length.out = 100),
  c_grandchildren = c(-1, 0, 1),
  Age             = mean(data_women$Age, na.rm = TRUE)
)

# predict() for clm returns a matrix of probabilities per category
probs <- predict(model_clm, newdata = newdat_clm, type = "prob")$fit
colnames(probs) <- levels(data_women$MOCA_cat)
```

```r
newdat_clm |>
  bind_cols(as.data.frame(probs)) |>
  pivot_longer(
    cols = all_of(levels(data_women$MOCA_cat)),
    names_to = "Category", values_to = "Probability"
  ) |>
  mutate(
    Category = factor(Category, levels = c("Severe", "Mild", "Normal")),
    Grandchildren = factor(c_grandchildren,
      levels = c(-1, 0, 1),
      labels = c(
        "Few grandchildren (-1 SD)",
        "Average grandchildren (0)",
        "Many grandchildren (+1 SD)"
      )
    )
  ) |>
  ggplot(aes(x = c_children, y = Probability, colour = Grandchildren)) +
  geom_line(linewidth = 1.1) +
  facet_wrap(~Category, ncol = 3) +
  scale_colour_manual(values = c("#2166ac", "#f4a582", "#1a7837")) +
  labs(
    x      = "Number of children (centred and scaled)",
    y      = "Predicted probability",
    colour = NULL
  ) +
  theme_minimal(base_size = 11) +
  theme(
    legend.position = "bottom",
    strip.text = element_text(face = "bold")
  )
```

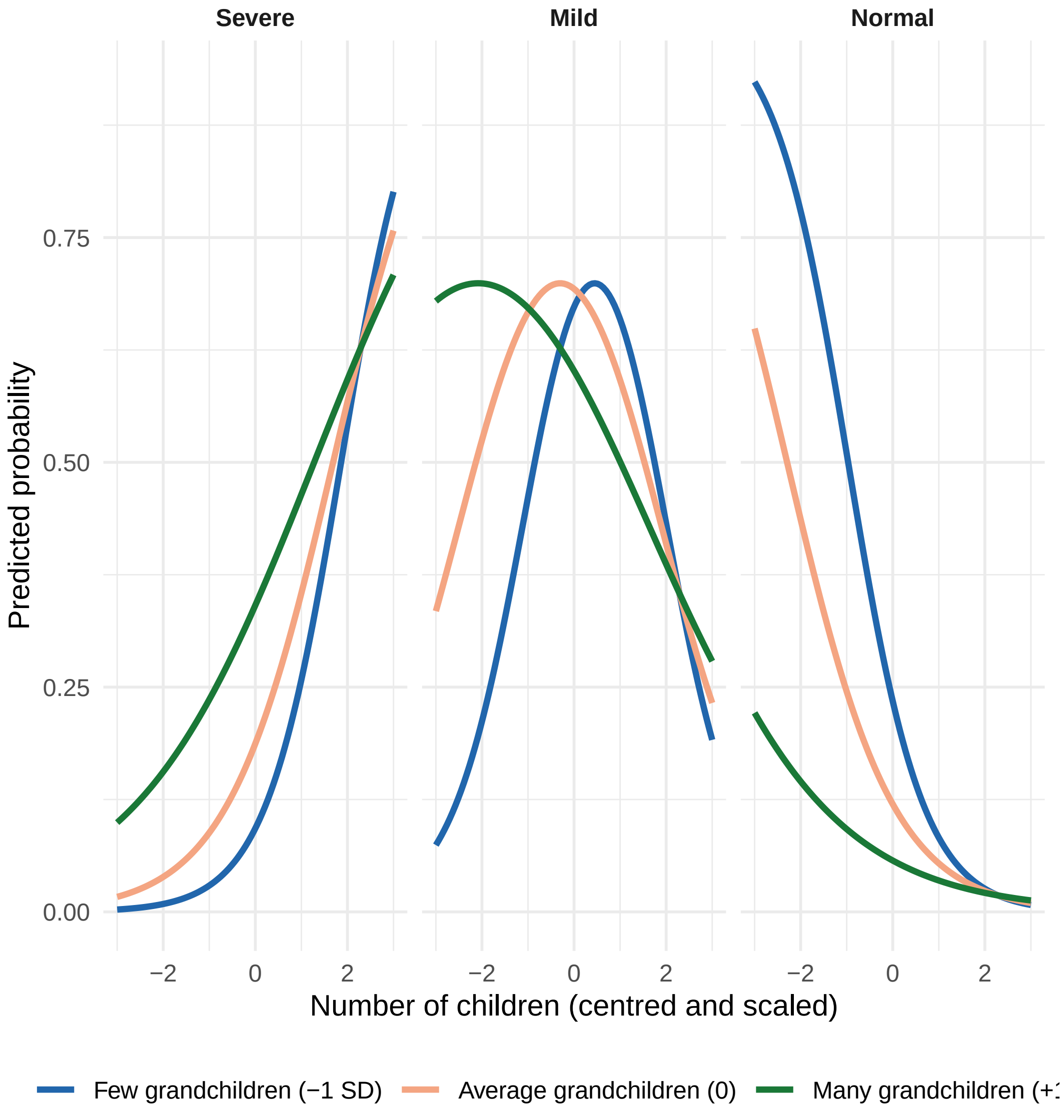

**Figure S4.** Predicted probabilities of each cognitive status category from the cumulative ordinal model, as a function of number of children (centred), at three levels of number of grandchildren and at mean age.

---

### 6 Model comparison

```
aic_vals <- tibble(
  Model = c(
    "Linear regression", "Poisson regression",
    "Cumulative ordinal logistic regression"
  ),
```

```
  AIC = c(AIC(model_lm), AIC(model_pois), AIC(model_clm))
) |>
  arrange(AIC) |>
  mutate(
    `ΔAIC` = round(AIC - min(AIC), 2),
    AIC    = round(AIC, 2)
  )

aic_vals |>
  flextable() |>
  autofit() |>
  theme_booktabs() |>
  add_footer_lines(
    "Note. ΔAIC = difference from the model with the lowest AIC."
  )
```

**Table S5.** Comparison of the three candidate models using the Akaike Information Criterion (AIC). Lower AIC indicates better fit; ΔAIC is computed relative to the best model.

| Model | AIC | ΔAIC |
|---|---|---|
| Cumulative ordinal logistic regression | 243.10 | 0.00 |
| Linear regression | 828.87 | 585.77 |
| Poisson regression | 841.82 | 598.72 |

Note. ΔAIC = difference from the model with the lowest AIC.

The cumulative ordinal model showed a substantially lower AIC than either the linear or Poisson models ($\Delta$AIC $>$ 500 for both), indicating a markedly better fit. This is expected given that the outcome is ordinal rather than continuous, and the ordinal model directly respects the measurement scale of the MOCA-Col categories. Accordingly, the ordinal model is used as the basis for inference in the manuscript.

---

### 7 Detailed results: Cumulative ordinal model

#### 7.1 Proportional odds assumption (Brant test)

```r
# polr refit required by the brant package
model_polr <- polr(
  MOCA_cat ~ Age + c_children * c_grandchildren,
  data   = data_women,
  method = "logistic"
)

brant(model_polr)
```

```
------------------------------------------------------------
Test for            X2  df  probability
------------------------------------------------------------
Omnibus             6.49    4   0.17
Age             3.78    1   0.05
c_children          0.58    1   0.44
c_grandchildren     1.58    1   0.21
c_children:c_grandchildren  0.69    1   0.41
------------------------------------------------------------

H0: Parallel Regression Assumption holds
```

The Brant test showed no evidence of violation of the proportional odds assumption (global p > .05), supporting the use of the cumulative model.

#### 7.2 Nominal effects

The `nominal_test()` function assesses whether the regression coefficients are constant across the cumulative logits (i.e., whether the proportional odds assumption holds for each predictor individually). A significant result indicates that the effect of a predictor varies across the thresholds.

```r
nominal_test(model_clm)
```

```
Tests of nominal effects

formula: MOCA_cat ~ Age + c_children * c_grandchildren
                           Df  logLik    AIC    LRT Pr(>Chi)
<none>                        -115.55 243.10
Age
c_children                  1 -114.12 242.24 2.8578  0.09093 .
c_grandchildren             1 -113.25 240.50 4.5928  0.03211 *
c_children:c_grandchildren  1 -114.84 243.68 1.4125  0.23464
---
```

```
Signif. codes:  0 '***' 0.001 '**' 0.01 '*' 0.05 '.' 0.1 ' ' 1
```

The nominal test indicated a significant effect for `c_grandchildren` ($p$ = .032), suggesting that its effect may not be strictly proportional across all thresholds. No significant nominal effects were found for `c_children` ($p$ = .091), the interaction term, or age. However, the Brant test showed no global violation of the proportional odds assumption, and this isolated result should be interpreted with caution given the limited statistical power to detect threshold-specific differences in a sample of this size. The proportional odds model is retained as a parsimonious and well-supported specification.

##### 7.3 Scale effects

The `scale_test()` function assesses whether any predictor influences the scale (dispersion) parameter of the model, analogous to heteroscedasticity in linear regression.

```
scale_test(model_clm)
```

```
Tests of scale effects

formula: MOCA_cat ~ Age + c_children * c_grandchildren
                           Df  logLik    AIC     LRT Pr(>Chi)
<none>                        -115.55 243.10
Age                         1 -115.46 244.91 0.18428   0.6677
c_children                  1 -114.67 243.34 1.75985   0.1846
c_grandchildren             1 -114.99 243.99 1.10708   0.2927
c_children:c_grandchildren  1 -115.22 244.43 0.66623   0.4144
```

Likelihood ratio tests indicated no evidence that any predictor influenced the scale parameter of the model (all $p > .10$), further supporting the adequacy of the proportional-odds specification.

##### 7.4 Collinearity (VIF)

```
# Refit as Poisson (same linear predictor) to use vif
model_vif <- glm(
  MOCACol ~ Age + c_children * c_grandchildren,
  family = poisson, data = data_women
)

vif(model_vif)
```

```
                       Age                 c_children
                  1.033486                   2.852256
           c_grandchildren c_children:c_grandchildren
```

```
                  3.739175                   2.586417
```

All VIF values are below 4 after centring and scaling, indicating negligible collinearity.

##### 7.5 Simple slopes

To decompose the interaction, we calculated the simple slope of number of children on cognitive status at three levels of number of grandchildren (-1, 0, and +1 SD).

```
slopes_clm <- emtrends(
  model_clm,
  ~c_grandchildren,
  var = "c_children",
  at  = list(c_grandchildren = c(-1, 0, 1))
)

summary(slopes_clm, infer = TRUE) |>
  as.data.frame() |>
  mutate(
    c_grandchildren = factor(c_grandchildren,
      levels = c(-1, 0, 1),
      labels = c(
        "Few grandchildren (-1 SD)",
        "Average grandchildren (0)",
        "Many grandchildren (+1 SD)"
      )
    ),
    c_children.trend = round(c_children.trend, 3),
    SE = round(SE, 3),
    lower.CL = round(asymp.LCL, 3),
    upper.CL = round(asymp.UCL, 3),
    z.ratio = round(z.ratio, 2),
    p.value = ifelse(p.value < .001, "< .001",
      sub("^0", "", sprintf("%.3f", p.value))
    )
  ) |>
  dplyr::select(
    c_grandchildren, c_children.trend, SE,
    lower.CL, upper.CL, z.ratio, p.value
  ) |>
  rename(
    `Level of grandchildren` = c_grandchildren,
```

```
    `Slope (children)`        = c_children.trend,
    `95% CI lower`            = lower.CL,
    `95% CI upper`            = upper.CL,
    z                         = z.ratio,
    p                         = p.value
  ) |>
  flextable() |>
  theme_booktabs() |>
  fontsize(size = 9, part = "all") |>
  fit_to_width(max_width = 6.5) |>
  add_footer_lines(
    "Note. Slopes represent the estimated effect of a one-SD increase in number
    of children on the log-odds of being in a higher cognitive category."
  )
```

**Table S6.** Simple slopes of the effect of number of children on cognitive status (MOCA-Col) at three levels of number of grandchildren.

| Level of grandchildren | Slope (children) | SE | 95% CI lower | 95% CI upper | z p |
|---|---|---|---|---|---|
| Few grandchildren (-1 SD) | -1.223 | 0.344 | -1.898 | -0.549 | -3.56 < .001 |
| Average grandchildren (0) | -0.869 | 0.305 | -1.467 | -0.272 | -2.85 .004 |
| Many grandchildren (+1 SD) | -0.516 | 0.291 | -1.085 | 0.054 | -1.77 .076 |

Note. Slopes represent the estimated effect of a one-SD increase in number
of children on the log-odds of being in a higher cognitive category.

```
# Pairwise contrasts between slopes
contrast(slopes_clm, method = "pairwise", adjust = "bonferroni")
```

```
 contrast                                  estimate     SE  df z.ratio p.value
 (c_grandchildren-1) - c_grandchildren0      -0.354 0.0927 Inf  -3.818  0.0004
 (c_grandchildren-1) - c_grandchildren1      -0.708 0.1850 Inf  -3.818  0.0004
 c_grandchildren0 - c_grandchildren1         -0.354 0.0927 Inf  -3.818  0.0004

P value adjustment: bonferroni method for 3 tests
```

The effect of having more children was significantly more negative at low levels of grandchildren than at high levels, confirming the moderation.

##### 7.6 Interaction plot

```
slopes_df <- summary(slopes_clm, infer = TRUE) |>
  as.data.frame() |>
  mutate(
    Grandchildren = factor(c_grandchildren,
      levels = c(-1, 0, 1),
      labels = c(
        "Few grandchildren (-1 SD)",
        "Average grandchildren (0)",
        "Many grandchildren (+1 SD)"
      )
    )
  )

ggplot(
  slopes_df,
  aes(
    x = Grandchildren, y = c_children.trend,
    ymin = asymp.LCL, ymax = asymp.UCL, colour = Grandchildren
  )
) +
  geom_hline(yintercept = 0, linetype = "dashed", colour = "grey50") +
  geom_errorbar(width = 0.15, linewidth = 0.8) +
  geom_point(size = 3) +
  scale_colour_manual(values = c("#2166ac", "#f4a582", "#1a7837")) +
  labs(
    x = "Level of grandchildren",
    y = "Slope of number of children\non log-odds of better\ncognitive status"
  ) +
  theme_minimal(base_size = 12) +
```

```r
  theme(
  legend.position = "none",
  axis.text.x = element_text(angle = 15, hjust = 1)
)
```

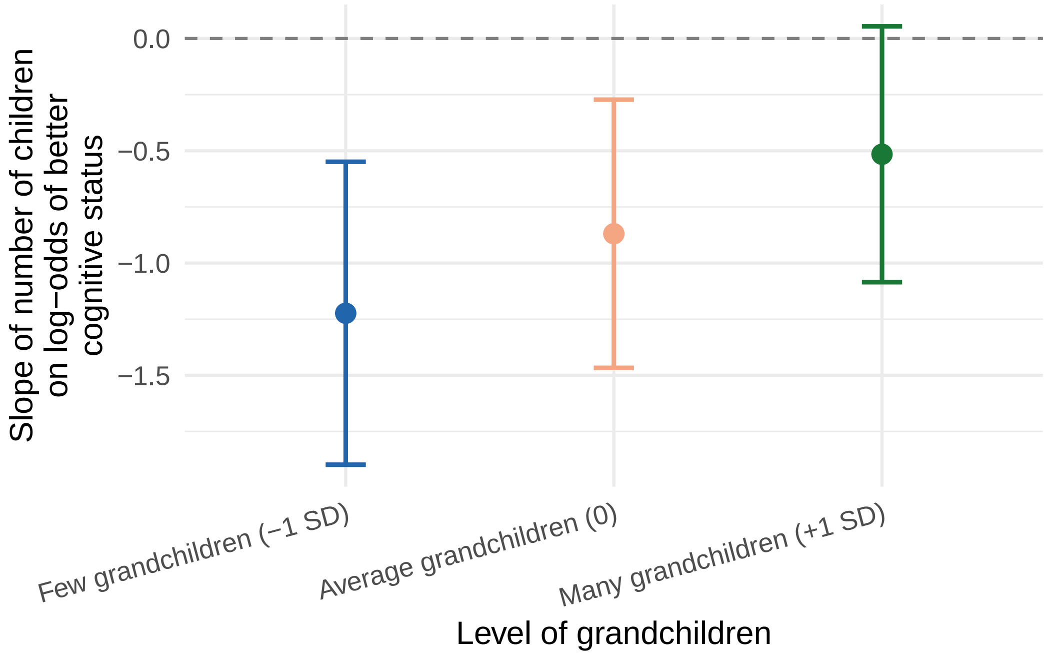

**Figure S5.** Simple slopes of the effect of number of children on the log-odds of better cognitive status, at three levels of number of grandchildren. Shaded bands represent 95% confidence intervals.

#### 7.7 Johnson–Neyman analysis

The Johnson–Neyman procedure identifies the precise range of the moderator (number of grandchildren) for which the focal effect (number of children) is or is not statistically significant at = .05.

```r
jn <- johnson_neyman(
  model  = model_clm,
  pred   = c_children,
  modx   = c_grandchildren,
  alpha  = .05
)

print(jn$bounds)
```

```
    Lower    Higher
0.8309587 5.1579800
```

```
jn$plot
```

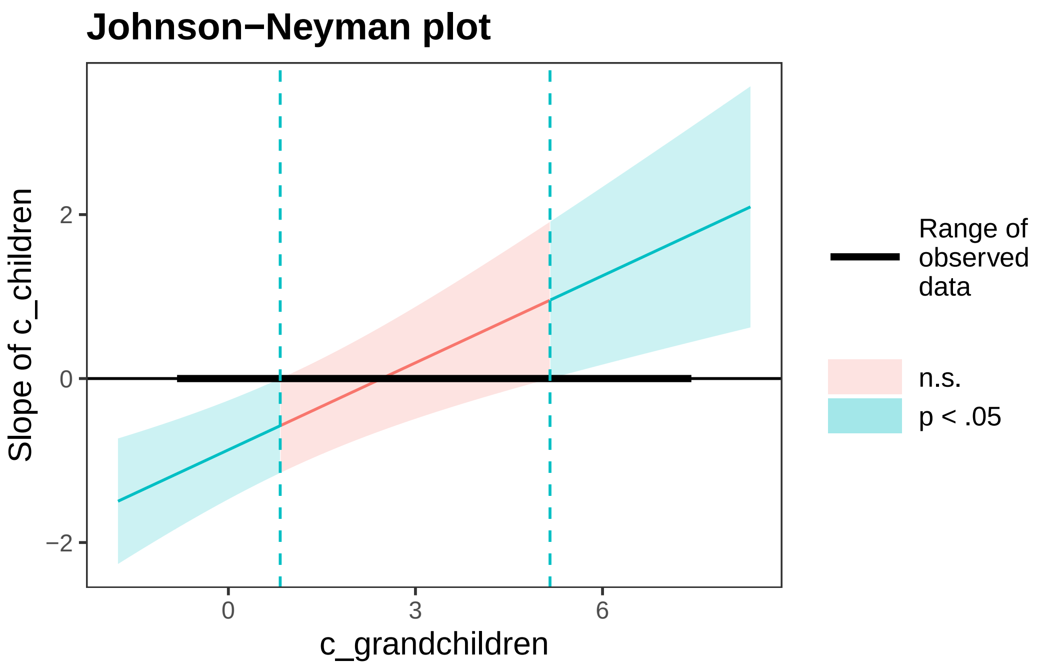

**Figure S6.** Johnson–Neyman plot. The shaded region indicates values of the number of grandchildren (centred and scaled) for which the effect of number of children on cognitive status is not statistically significant (p > .05). Vertical dashed lines mark the significance boundaries.

The Johnson–Neyman analysis showed that the effect of number of children on cognitive status was statistically significant only at relatively low values of number of grandchildren. Beyond the upper boundary, the association attenuated and ceased to be significant, indicating that the presence of more grandchildren progressively moderates the negative effect of number of children on cognition.

---

### 8 Manuscript figure

The following figure combines the two most informative results of the interaction analysis into a single two-panel display intended for inclusion in the manuscript.

```
library(patchwork)

# --- Panel A: Johnson-Neyman ---
p_jn <- jn$plot +
  labs(
    title = "A",
    x     = "Number of grandchildren (centred and scaled)",
```

```
    y     = "Slope of number of children"
  ) +
  theme_minimal(base_size = 10) +
  guides(fill = guide_legend(title = "Significance")) +
  theme(
    plot.title      = element_text(face = "bold", size = 14),
    legend.position = "bottom"
  )

# --- Panel B: Simple slopes ---
p_slopes <- slopes_df |>
  mutate(
    Grandchildren = factor(c_grandchildren,
      levels = c(-1, 0, 1),
      labels = c("Few\n(-1 SD)", "Average\n(0)", "Many\n(+1 SD)")
    )
  ) |>
  ggplot(aes(
    x      = Grandchildren, y = c_children.trend,
    ymin   = asymp.LCL, ymax = asymp.UCL,
    colour = Grandchildren
  )) +
  geom_hline(yintercept = 0, linetype = "dashed", colour = "grey50") +
  geom_errorbar(width = 0.12, linewidth = 0.8) +
  geom_point(size = 3.5) +
  scale_colour_manual(values = c("#2166ac", "#f4a582", "#1a7837")) +
  labs(
    title = "B",
    x     = "Number of grandchildren",
    y     = "Slope of number of children\n(log-odds of better cognitive status)"
  ) +
  theme_minimal(base_size = 10) +
  theme(
    plot.title      = element_text(face = "bold", size = 14),
    legend.position = "none"
  )

# --- Panel C: Predicted probabilities by MOCA category ---
p_emmip <- emmip(
```

```
  model_clm,
  MOCA_cat ~ c_children | c_grandchildren,
  mode = "prob",
  CIs   = TRUE,
  at    = list(
    c_children      = seq(-1, 1, length.out = 50),
    c_grandchildren = c(-1, 0, 1)
  ),
  plotit = FALSE
) |>
  mutate(
    c_grandchildren = factor(c_grandchildren,
      levels = c(-1, 0, 1),
      labels = c("Few grandchildren (-1 SD)",
                 "Average grandchildren (0)",
                 "Many grandchildren (+1 SD)")
    )
  ) |>
  ggplot(aes(x = xvar, y = yvar, colour = tvar, fill = tvar)) +
  geom_ribbon(aes(ymin = LCL, ymax = UCL), alpha = 0.15, colour = NA) +
  geom_line(linewidth = 1) +
  facet_wrap(~ c_grandchildren) +
  scale_colour_manual(
    values = c("#d62728", "#ff7f0e", "#2ca02c"),
    name   = "Cognitive status"
  ) +
  scale_fill_manual(
    values = c("#d62728", "#ff7f0e", "#2ca02c"),
    name   = "Cognitive status"
  ) +
  labs(
    title = "C",
    x     = "Number of children (centred and scaled)",
    y     = "Predicted probability"
  ) +
  theme_minimal(base_size = 10) +
  theme(
    plot.title      = element_text(face = "bold", size = 14),
    legend.position = "bottom",
```

```
    strip.text      = element_text(face = "bold")
  )

# --- Final figure ---
fig1 <- (p_jn + p_slopes) /
  p_emmip + plot_layout(heights = c(1, 1.2))
fig1
```

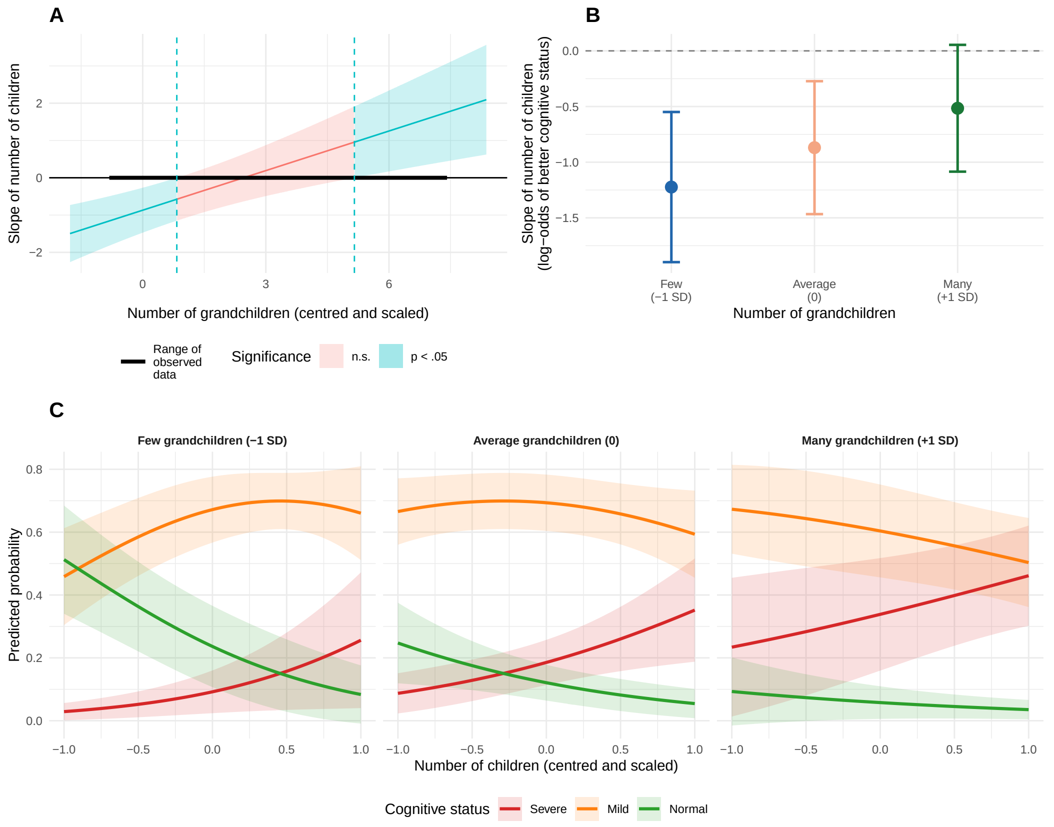

**Figure 1.** Interaction between number of children and number of grandchildren on cognitive status. (A) Johnson–Neyman plot showing the range of number of grandchildren (centred and scaled) for which the association with number of children is statistically significant (blue shading, $p < .05$) or not (red shading, $p > .05$). Dashed vertical lines mark the significance boundaries. (B) Simple slopes of the effect of number of children on the log-odds of better cognitive status at three levels of number of grandchildren. Points represent slope estimates; error bars represent 95% confidence intervals. The dashed horizontal line indicates a slope of zero. (C) Predicted probabilities of each cognitive status category (Severe, Mild, Normal) as a function of number of children (centred and scaled), separately for low (-1 SD), average (0), and high (+1 SD) levels of number of grandchildren.

### 9 Session information

```
toLatex(sessionInfo(), locale = FALSE)
```

- R version 4.5.3 (2026-03-11), `x86_64-pc-linux-gnu`
- Running under: `CachyOS`
- Matrix products: default
- BLAS: `/usr/lib/libblas.so.3.12.0`
- LAPACK: `/usr/lib/liblapack.so.3.12.0`
- Base packages: base, datasets, graphics, grDevices, methods, stats, utils
- Other packages: AER 1.2-16, brant 0.3-0, broom 1.0.12, car 3.1-5, carData 3.0-6, dplyr 1.2.1, emmeans 2.0.3, flextable 0.9.11, forcats 1.0.1, ggplot2 4.0.2, interactions 1.2.0, lmtest 0.9-40, lubridate 1.9.5, MASS 7.3-65, ordinal 2025.12-29, patchwork 1.3.2, purrr 1.2.1, readr 2.2.0, sandwich 3.1-1, stringr 1.6.0, survival 3.8-6, tibble 3.3.1, tidyr 1.3.2, tidyverse 2.0.0, zoo 1.8-15
- Loaded via a namespace (and not attached): abind 1.4-8, askpass 1.2.1, backports 1.5.1, bit 4.6.0, bit64 4.6.0-1, broom.mixed 0.2.9.7, cli 3.6.6, coda 0.19-4.1, codetools 0.2-20, compiler 4.5.3, crayon 1.5.3, data.table 1.18.2.1, digest 0.6.39, estimability 1.5.1, evaluate 1.0.5, farver 2.1.2, fastmap 1.2.0, fontBitstreamVera 0.1.1, fontLiberation 0.1.0, fontquiver 0.2.1, Formula 1.2-5, furrr 0.4.0, future 1.70.0, gdtools 0.5.0, generics 0.1.4, globals 0.19.1, glue 1.8.0, grid 4.5.3, gtable 0.3.6, hms 1.1.4, htmltools 0.5.9, jsonlite 2.0.0, jtools 2.3.1, knitr 1.51, labeling 0.4.3, lattice 0.22-9, lifecycle 1.0.5, listenv 0.10.1, magrittr 2.0.5, Matrix 1.7-5, multcomp 1.4-30, mvtnorm 1.3-6, nlme 3.1-169, numDeriv 2016.8-1.1, officer 0.7.3, openssl 2.3.5, otel 0.2.0, pander 0.6.6, parallel 4.5.3, parallelly 1.46.1, pillar 1.11.1, pkgconfig 2.0.3, R6 2.6.1, ragg 1.5.2, RColorBrewer 1.1-3, Rcpp 1.1.1, rlang 1.2.0, rmarkdown 2.31, rstudioapi 0.18.0, S7 0.2.1, scales 1.4.0, splines 4.5.3, stringi 1.8.7, systemfonts 1.3.2, textshaping 1.0.5, TH.data 1.1-5, tidyselect 1.2.1, timechange 0.4.0, tools 4.5.3, tzdb 0.5.0, ucminf 1.2.3, uuid 1.2-2, vctrs 0.7.2, vroom 1.7.1, withr 3.0.2, xfun 0.57, xml2 1.5.2, xtable 1.8-8, yaml 2.3.12, zip 2.3.3
